# Phospho-Tune: Enhanced Structural Modeling of Phosphorylated Protein Interactions

**DOI:** 10.1101/2024.02.29.582580

**Authors:** Ernest Glukhov, Veranika Averkava, Sergei Kotelnikov, Darya Stepanenko, Thu Nguyen, Julie C. Mitchell, Carlos Simmerling, Sandor Vajda, Andrew Emili, Dzmitry Padhorny, Dima Kozakov

**Affiliations:** Department of Applied Mathematics and Statistics, Stony Brook University, Stony Brook, 11794, NY, USA; Department of Computer Science, Stony Brook University, Stony Brook, 11794, NY, USA; Department of Chemistry, Stony Brook University, Stony Brook, 11794, NY, USA; Laufer Center for Physical and Quantitative Biology, Stony Brook University, Stony Brook, 11794, NY, USA; Biosciences Division, Oak Ridge National Laboratory, Oak Ridge, TN, USA; Department of Biomedical Engineering, Boston University, Boston, 02215, MA, USA; Center for Network Systems Biology, Boston University, Boston, MA, USA; Department of Biochemistry, Boston University School of Medicine, Boston, MA, USA; Department of Biology, Boston University, Boston, MA, USA

**Keywords:** Phosphorylation, Protein-Peptide Interactions, Structural Prediction, Deep learning, AlphaFold, Fine-tuning, Cellular Signaling

## Abstract

In computational biology, accurate prediction of phosphopeptide-protein complex structures is essential for understanding cellular functions and advancing drug discovery and personalized medicine. While AlphaFold has significantly improved protein structure prediction, it faces accuracy challenges in predicting structures of complexes involving phosphopeptides possibly due to structural variations introduced by phosphorylation in the peptide component. Our study addresses this limitation by refining AlphaFold to improve its accuracy in modeling these complex structures. We employed weighted metrics for a comprehensive evaluation across various protein families. The enhanced model notably outperforms the original AlphaFold, showing a substantial increase in the weighted average local distance difference test (lDDT) scores for peptides: from 52.74 to 76.51 in the Top 1 model and from 56.32 to 77.91 in the Top 5 model. These advancements not only deepen our understanding of the role of phosphorylation in cellular signaling but also have extensive implications for biological research and the development of innovative therapies.

## Introduction

Phosphorylation, a critical post-translational modification, plays a pivotal role in regulating protein function and signaling pathways within cells [18]. This biochemical process involves the addition of a phosphate group to a protein or a peptide, typically mediated by enzymes such as kinases [14, 21]. Phosphorylation can cause significant changes in the protein’s structure impacting its interaction with other molecules, including peptides [6, 10]. These interactions are crucial for various cellular processes, including cell cycle regulation, growth, apoptosis, and immune responses [1, 13, 12, 16, 8].

The significance of phosphorylation in protein-peptide interactions lies in its ability to modulate the function of proteins [17]. It can activate or deactivate enzymes, alter subcellular localization, or promote or prevent interactions with other proteins and peptides. These modulatory effects are central to cellular signaling and are intricately linked to numerous physiological processes and disease states. For example, aberrant phosphorylation patterns are often associated with diseases like cancer, diabetes, and neurodegenerative disorders, making it a key area of interest in therapeutic research[11, 19, 4].

AlphaFold has revolutionized protein structure prediction through its utilization of deep learning [9]. It’s notably effective in determining protein structures, often outperforming traditional methods. However, its capability in accurately predicting phosphopeptide-protein complexes is not as profound due to its training on a diverse protein dataset, which, while beneficial for general applications, might not capture the intricate details required for phosphopeptide-protein interactions. To address this, our research fine-tunes AlphaFold using a curated dataset enriched with high-quality phosphopeptide-protein complex data. By tailoring the model to this specific domain, we aim to substantially enhance its precision in this critical area.

In this study, we introduce the Phospho-Tune model, an AlphaFold-based approach, specifically fine-tuned to enhance the accuracy in modeling phosphopeptide-protein complexes. Our refined model not only provides better structural predictions but also significantly improves the confidence assessment of these predictions, a crucial aspect for practical applications.

The overall pipeline of predicting the three-dimensional structures of phosphopeptide-protein complexes using the proposed Phospho-Tune model is plotted in the Figure 5.

## Methodology

### Data collection and preprocessing

To specifically tailor our model for the structural prediction of phosphopeptide-protein complexes, we constructed a specialized dataset sourced from the Protein Data Bank (PDB) [2]. Our selection criteria involved identifying samples that contained phosphorylated residues of types SEP (phosphoserine), TPO (phosphothreonine), and PTR (phosphotyrosine). In addition to these criteria, we also filtered out samples with non-standard amino acids and a resolution greater than 3 angstroms to ensure high-quality, reliable data. Furthermore, we excluded any cases where the phosphorylated residue was situated more than 5 angstroms away from the nearest contacted protein chain to ensure relevance in our dataset.

The AlphaFold model uses Multiple Sequence Alignments (MSAs) as a key input feature to predict protein structures. In our study, we also adapted this approach for peptide analysis. For certain peptides, when a corresponding UniProt ID was identifiable, we generated MSAs for the UniProt sequences using MMseqs2. From these alignments, we aligned the UniProt sequence with the peptide sequence and extracted segments specifically relevant to the peptides. To ensure a targeted focus on local structural environments pertinent to phosphorylation events, we cropped all peptides to include only the region encompassing 4 residues upstream and downstream of the phosphorylated residue. By concentrating on these specific areas, we aim to capture spatial relationships and interactions critical to understanding the molecular mechanisms underlying phosphorylation events.

The final dataset comprises 967 structures, and to ensure our model’s adaptability across diverse protein families and structural configurations, we conducted a redundancy analysis using a clustering approach. We used HHsearch [20] against the PDB100 database, generating hhr files that identify distant homologies. We then clustered these samples based on high scores in the ‘Prob’ (probability) column of the hhr files, grouping together similar proteins. This method allowed us to distill our dataset, ensuring a non-redundant and clean test set.

Our dataset was further divided into three subsets for training, validation, and testing. The training set, consisting of 512 samples, was organized chronologically based on release dates, spanning from 1993-10-31 to 2017-05-24. The remaining samples were then divided into the validation and test sets, considering clusters. The test set includes 228 samples, and the validation set comprises 227 samples. The training set and the validation set were used for fine-tuning and hyperparameter optimization, while the test set was reserved for the final model evaluation.

### Model performance evaluation

In our study, we used Root Mean Square Deviation (RMSD), Local Distance Difference Test (lDDT) [15], and predicted Local Distance Difference Test (plDDT) as key metrics to evaluate the structural accuracy. RMSD quantifies the average distance between the atoms of superimposed proteins, offering a measure of the similarity between protein structures. Lower RMSD values indicate greater structural similarity. However, RMSD can be sensitive to outliers and does not account for local structural variations. lDDT addresses this by assessing the local conformational differences between structures, considering the distances between all atom pairs within defined cutoffs, thereby providing a more nuanced view of local structural accuracy and stability. plDDT, an extension used in models like AlphaFold, predicts the lDDT score for each residue in a protein structure, offering a residue-level confidence measure of the predicted model’s accuracy. The combination of these metrics allows for a comprehensive and detailed evaluation of the structural predictions, ensuring both global and local structural features are accurately captured and assessed in our study.

In addition, to assess how well our model predicts important parts of the complex structure, such as peptides and phosphorylated residues, we used region-specific masks. By applying them, we calculated our key metrics such as alpha-carbon peptide RMSD (C*α*-pRMSD) and alpha-carbon RMSD for the phosphorylated residue (C*α*-phosRMSD), as well as masked lDDT and plDDT metrics to measure accuracy only in these specific areas within the predicted structure.

### Model implementation

In our current research, we advance the custom implementation of AlphaFold model introduced in our previous paper [7], focusing on enhanced flexibility and efficiency. We have optimized memory usage through checkpointing and efficient resource management across multi-GPU and multi-node setups using PyTorch-Lightning [5]. Additionally, the incorporation of Weights & Biases [3] has allowed us to establish a robust pipeline for the model’s development, experimentation, and reporting. These improvements collectively contribute to a more systematic and scalable framework for protein structure prediction.

#### Phosphorylation integration in model’s architecture

To highlight the significance of phosphorylation in our specific task, we implemented targeted architectural modifications in AlphaFold aimed at enhancing the model’s understanding of their positions within the peptide sequence. These adjustments are crafted to strengthen the model’s capacity to recognize and interpret the spatial characteristics associated with phosphorylation.

Specifically, three new layers were introduced to the InputEmbedding module. These layers are designed for preprocessing and incorporating phosphorylation information, which are indicated by a masked feature revealing the positions of phosphorylated residues in the peptide sequence. These new layers integrate phosphorylation information into both amino acid and pair representations. The first layer, employing a linear transformation, interprets phosphorylation features to capture relationships with amino acid sequences, while the other two contribute phosphorylation-specific features to pair representations, providing the model with insights into how phosphorylation influences interactions between amino acids.

#### Enhancements in PredictedLDDT Module

In an effort to enhance the reliability of plDDT scores, which are crucial for assessing the quality of protein structure predictions, we have refined the PredictedLDDT module. The original module, consisting of two linear layers, was expanded to include additional linear layers, each paired with a Rectified Linear Unit (ReLU) activation function. This adjustment allows for variable layer configurations and introduces the number of layers as a tunable parameter during the fine-tuning process.

In addition, our implementation introduces a new approach for the separate fine-tuning of the PredictedLDDT module within the AlphaFold architecture. This process involves isolating the module, separate fine-tuning, and then integrating the refined weights back into the full AlphaFold model. This independent fine-tuning of the PredictedLDDT module is particularly advantageous, as it enhances plDDT scores efficiently without extensive computational resources. This targeted approach optimizes plDDT predictions for specific applications, eliminating the need for complete AlphaFold model retraining.

### Experiments

In this section, we present a series of experiments that systematically evaluate important aspects of our study. These experiments explore key components such as data preprocessing techniques, the model’s architecture, and our training methodology. Our goal is to provide a detailed analysis of each element, showcasing their individual and combined impact on the overall performance of our system.

#### Incorporating peptide MSAs

In our study, we examined how incorporating Peptide Multiple Sequence Alignments, which highlight patterns of similarity and difference in peptide sequences, could enhance model accuracy in predicting the structures of phosphorylated peptide-protein interactions. We conducted training experiments both with and without using these alignments as the model’s input features to assess their impact on predictive performance.

#### AlphaFold multimer v.2.2 vs. v.2.3

In our experiments, we fine-tuned two different versions of AlphaFold multimer, versions 2.2 and 2.3. This approach was designed to understand and analyze the differences between these iterations and to evaluate their respective strengths and capabilities. Notably, version 2.3 has minor yet significant architectural enhancements compared to version 2.2. Moreover, it can process a greater number of sequences from Multiple Sequence Alignments.

#### Additional Evoformer Blocks

In our experiments, we explored modifications of the AlphaFold architecture by incorporating additional layers known as Evoformer blocks. We added from one to ten extra blocks to assess their impact on the model’s performance. The objective was to determine whether increasing the model’s complexity improved its ability to accurately predict complex structures.

#### PredictedLDDT Module Architecture

In the ‘Model Implementation’ section, under the ‘PredictedLDDT Module’ subsection, we describe enhancements made to the PredictedLDDT module aimed at improving the accuracy of predicted lDDT scores. We experimented with adding between 1 to 8 additional linear layers, each paired with a Rectified Linear Unit (ReLU), to the module’s original two-layer architecture.

#### Focused Training Methods

In our experimental setup, we explored the optimization of model performance through the adjustment of weights assigned to predictions concerning peptide structures. This was accomplished by modifying the weights in a loss function specifically designed for peptide-focused part of the predicted structure. We tested a range of weight values, extending from 0.0, which implies no additional loss, to 5.0, signifying an increase of the initial loss by a factor of five for the peptide part. Furthermore, we implemented and evaluated a preprocessing technique that involved masking out residues in peptide sequences where atom distances exceeded 5Å from the protein chain. By focusing on regions in close proximity to the protein chain, which are generally more biologically significant, the model could better prioritize and learn from the most pertinent structural features of the peptides.

#### Hyperparameter Optimization

In our efforts to improve the predictive accuracy of our model, a key focus was placed on hyperparameter tuning. We tested various hyperparameters, each playing a specific role in the model’s learning and performance.

- Learning Rate (LR): A fundamental parameter that determines the step size at each iteration during the model’s training, directly impacting the convergence speed and quality. We used values in range from 1e-3 to 1e-7.
- Learning Rate Schedulers: These tools adjust the learning rate during training, following specific patterns to improve convergence. We tested several schedulers including CyclicLR, OneCycleLR, StepLR, LinearLR, CosineAnnealingLR, and CosineAnnealingWarmRestarts.
- LR Gamma, LR Multiplier, LR Step Size Up, and LR Warm-up: These parameters are associated with learning rate adjustment, helping to fine-tune the model’s training process.
- Accumulate grad batches: This parameter allows for the accumulation of gradients over multiple batches, simulating a larger effective batch size, which can be crucial for model stability and performance.

## Results

### Phosphorylation integration in model’s architecture

Analyzing the predictive performance of the model before and after the integration of information about the positions of phosphorylated residues within the peptide sequence into the model’s architecture revealed a significant improvement in the model’s ability to predict the structure of phosphopeptides. As illustrated in Figure 1, using the representative sample (PDB ID: 7oqj), we can observe the practical implications of this modification. In cases where the peptide contains multiple serine residues, the initial model, lacking this modification, shifted the peptide structure within the binding site, causing it to twist. However, the updated model accurately identifies and positions the correct phosphorylated serine at its true location.

**Fig. 1:**
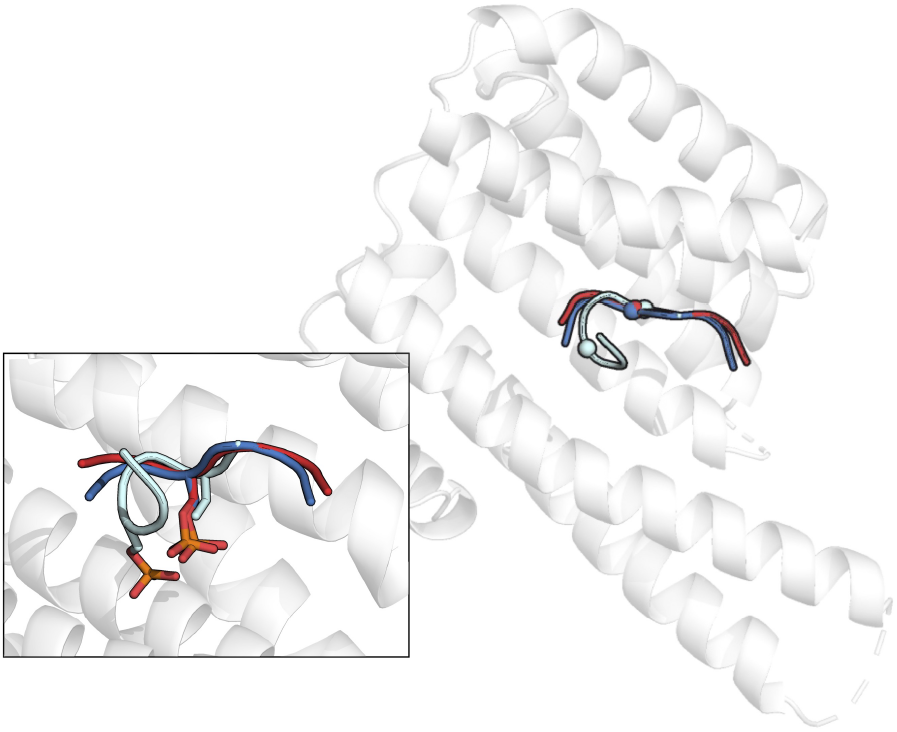
Model Comparison in Predicting Phosphorylated Residue Structure. The overall protein structure is shown with a zoomed view of peptide alignments: red indicates the true structure, cyan represents the original model’s prediction which places a wrong serine residue at the expected phosphorylated residue position, and blue depicts the updated model’s prediction. The updated model correctly identifies and aligns the phosphorylated serine, highlighting its enhanced predictive accuracy.

Additionally, for this sample, we conducted an experiment by relocating the position of the phosphorylated serine residue in a masked feature to a non-phosphorylated serine. Specifically, this serine was placed two positions to the left from the actual phosphorylated residue in the peptide sequence. The manipulation resulted in a misalignment in the model’s prediction, where it incorrectly identified a non-phosphorylated serine as the phosphorylated residue. This underscores the significance of incorporating phosphorylation information into the model to enhance its accuracy in modeling the structure of phosphopeptides.

Furthermore, the integration of phosphorylation information in the model led to faster convergence during training. This is demonstrated by consistently lower mean structure loss values for the peptide segment in Figure 2, where the average is calculated across five models. These results suggest that the model with phosphorylation integration achieves optimal predictive capabilities more efficiently, emphasizing the valuable impact of this architectural enhancement on both accuracy and efficiency in our task.

**Fig. 2:**
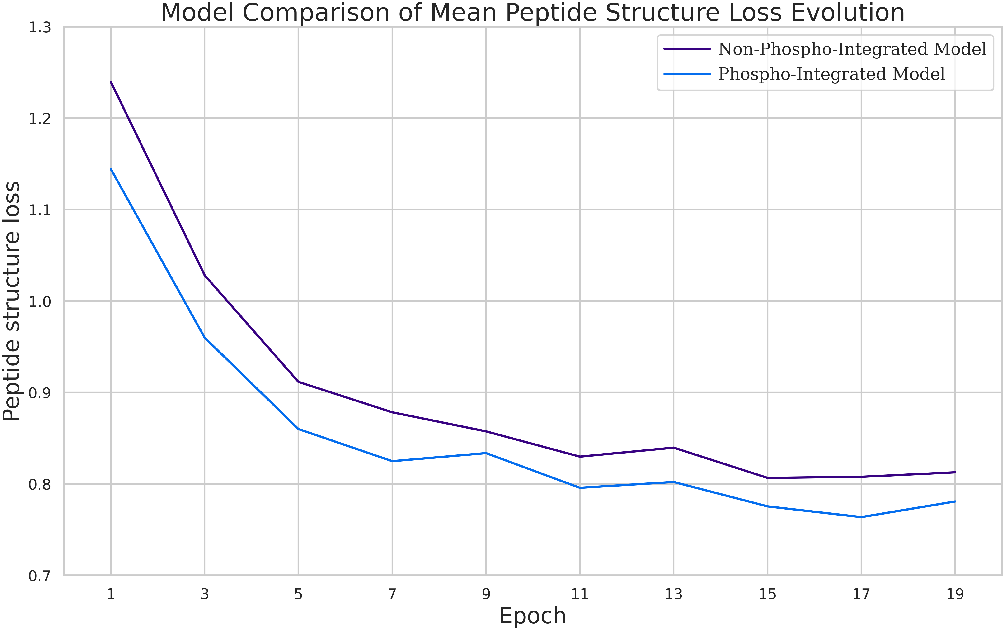
Comparison of mean peptide structure loss during multiple training epochs between Phosphorylation-Integrated and Non-Integrated models. The plot illustrates averaged loss values across five models, emphasizing accelerated training convergence in the Phosphorylation-Integrated model.

### PredictedLDDT module

The scatter plots in Figure 3 compare the mean predicted and actual lDDT scores for peptide residues using two models: the original AlphaFold and the same model but with the fine-tuned PredictedLDDT module. The dotted diagonal line represents the ideal prediction where predicted scores perfectly match the actual scores. The data points of the enhanced model are notably closer to this line than those of the baseline, indicating a more accurate prediction of lDDT scores and thus, more reliable protein structure predictions. This is supported by a significant reduction in the Mean Absolute Error (MAE) between true and predicted lDDT scores for peptides—from an average MAE of 15.06 for the original AlphaFold to an average of 7.8 in the model with the fine-tuned PredictedLDDT module. The ability to train the PredictedLDDT module separately for specific tasks and subsequently integrate it into the AlphaFold model underscores a significant advancement in our modeling approach, offering flexibility and task-specific optimization without the need to retrain the entire network.

**Fig. 3:**
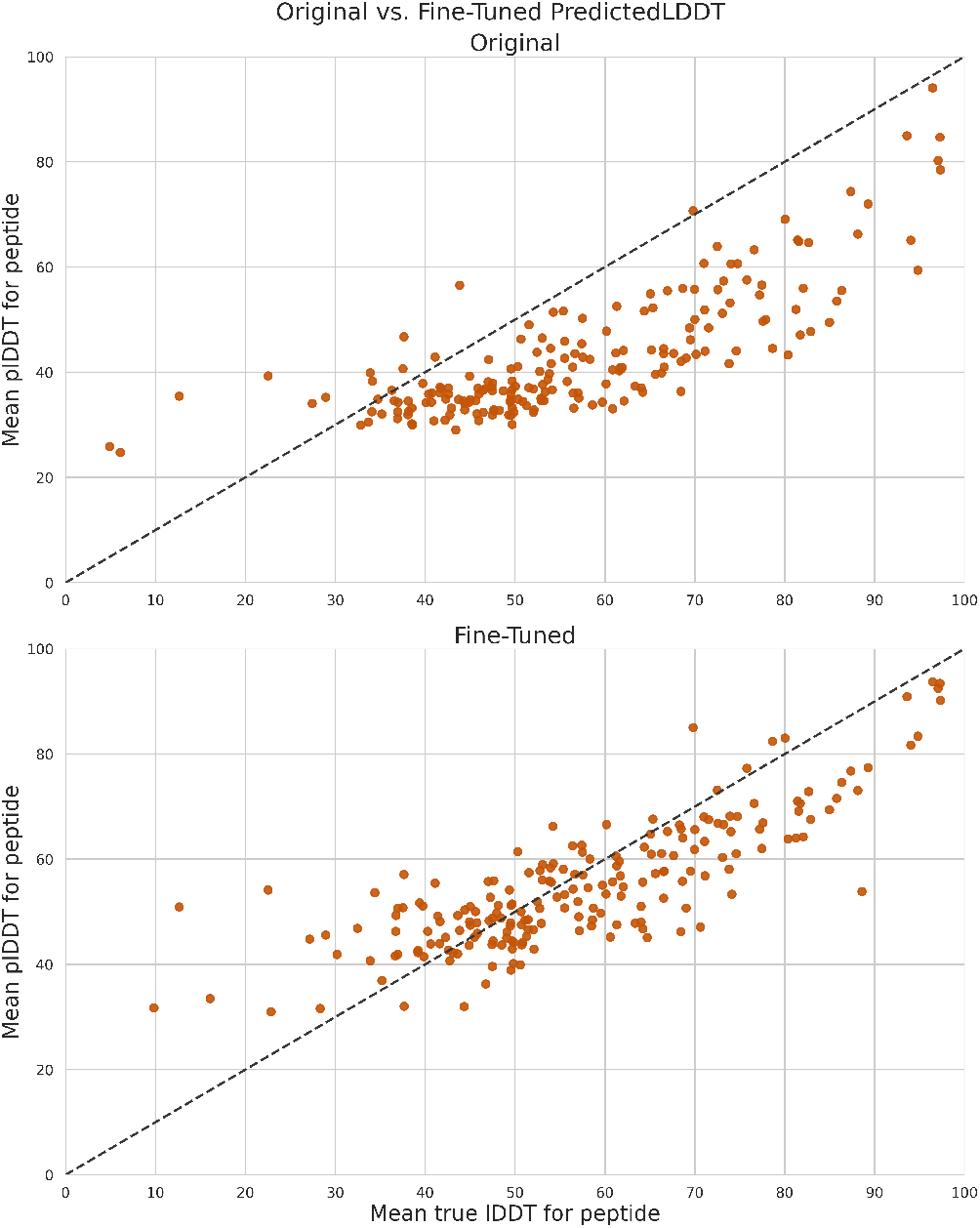
Scatter plots visually demonstrate the impact of separate fine-tuning by comparing the alignment between predicted lDDT scores and actual scores in the baseline model versus the baseline model featuring a fine-tuned PredictedLDDT module.

### Optimal Parameter Settings

Following an extensive experimentation process aimed at finding the optimal model’s architecture and training methodology, we have identified the best set of parameters for our specialized AlphaFold-based model:

#### Additional Evoformer Blocks

Experimenting with the number of additional Evoformer blocks, we discovered that increasing the original 48 blocks positively impacted the model’s performance. Specifically, for our task, incorporating 1 additional blocks proved to be the most effective. This configuration enhanced the model’s ability to capture intricate details of protein structures, showcasing an optimal balance achieved between model complexity and performance enhancement.

#### Focused Loss Function

Our experiments, which aimed to prioritize the importance assigned to the prediction of peptide structures within protein complexes through weighted adjustments to the loss function, did not result in substantial improvements in the overall model performance.

#### Hyperparameters

Among the hyperparameters explored, the learning rate proved to be the most critical factor influencing the model’s behavior and performance. We set the learning rate at 0.00001, noting that lower rates enhance training stability by allowing the model to take smaller optimization steps. Furthermore, we implemented the CyclicLR scheduler with a LR Gamma value of 0.9, a LR Multiplier value of 25, and a LR Step Size Up value of 2000. While the impact of these parameters on convergence speed and model stability was less pronounced than the learning rate itself, careful consideration of these parameters remains essential. We configured the accumulate grad batches parameter to 3. Since we used 3 GPUs for training, this configuration effectively translated to a real batch size of 9, allowing for more precise gradient updates and proving advantageous at this stage of the model’s training.

After fine-tuning the full AlphaFold multimer model, we focused on improving its lDDT score predictions by fine-tuning the PredictedLDDT module separately. In this process, we made adjustments to the module’s architecture and hyperparameters. Specifically, we used a OneCycleLR scheduler, added 8 additional linear layers, and set the learning rate to 0.05, with a LR multiplier of 2 and LR Warm-Up of 0.1. This allowed us to make our lDDT score predictions more accurate, thereby enhance the overall performance of our Top 1 model.

To assess the performance of both the original AlphaFold model and our fine-tuned version on our test set, we utilized all five multimer models available in AlphaFold. We assessed each sample in our test set by analyzing key metrics for the predicted structures produced by each model. The optimal result for each sample was selected based on plDDT scores for the Top 1 model, with preference given to the one with the highest plDDT score for the peptide part. For the Top 5 model, the selection was determined by the highest lDDT score for the peptide component obtained during the evaluation process.

During our initial fine-tuning with AlphaFold v.2.2, we achieved a 14.2% improvement for the Top 1 model and a 12.98% improvement for the Top 5 model in the weighted average lDDT score for peptides compared to their respective baselines. However, fine-tuning with version 2.3 significantly enhanced our predictive capabilities. In version 2.3, the fine-tuned Top 1 model surpassed the baseline by 31.07% in the weighted average lDDT score for peptides, while the Top 5 model exhibited a 27.71% improvement in the weighted average lDDT score for peptides. This version upgrade also led to more stable training, characterized by reduced fluctuations and smoother convergence patterns.

In Figure 4, we compare the performances of the fine-tuned AlphaFold multimer models v.2.3 (Top 1 and Top 5 models) with their original AlphaFold v.2.3 counterparts. This comparison is based on examining the weighted average lDDT scores specific to peptides. We apply a weighting scheme considering the diversity of protein families in our test set. This scheme assigns weights to samples based on their respective protein families’ sizes, with higher weights assigned to samples from less prevalent families. This ensures a balanced evaluation that emphasizes the contribution of smaller families in assessing the overall model performance. The comparison reveals that the fine-tuned models showcase enhanced predictive accuracy compared to their original AlphaFold counterparts.

**Fig. 4:**
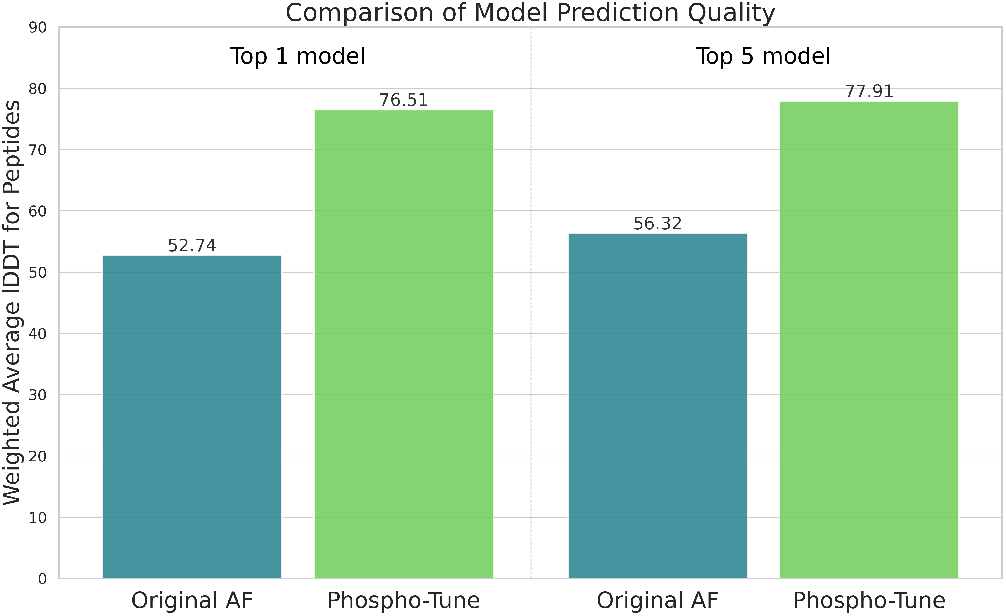
Comparison of Peptide Pose Prediction Accuracy between the Original and Fine-Tuned AlphaFold Models (v.2.3) for both Top 1 and Top 5 models. For the Top 1 model, we select predictions with the highest predicted lDDT scores among the five fine-tuned AlphaFold models, while for the Top 5 model, predictions are chosen based on the highest real lDDT scores within the same trained set of five models. The evaluation is based on the weighted average lDDT scores for peptides.

### Comparative Analysis

The comparison between our refined Phospho-Tune model and the baseline AlphaFold model is illustrated across 14 case studies representing various protein families (Examples A to N) in Figure 6. Phospho-Tune varies in prediction accuracy but consistently outperforms AlphaFold by more accurately modeling overall peptide structures and specifically phosphorylated residues. For instance, in Example B (PDB ID: 7nrk), Phospho-Tune achieves a significantly lower C*α*-pRMSD compared to AlphaFold (0.54 Å vs. 2.97 Å). In some cases, such as Example J (PDB ID: 6l03), Phospho-Tune demonstrates moderate accuracy in predicting peptide component of the complex structure. In other cases, like Example M (PDB ID: 6rh6), both models show prediction discrepancies, suggesting room for further improvement. Overall, Phospho-Tune demonstrates higher predictive accuracy for phosphopeptide-protein complex structures compared to AlphaFold, even in challenging scenarios encountered by both models.

**Fig. 5:**
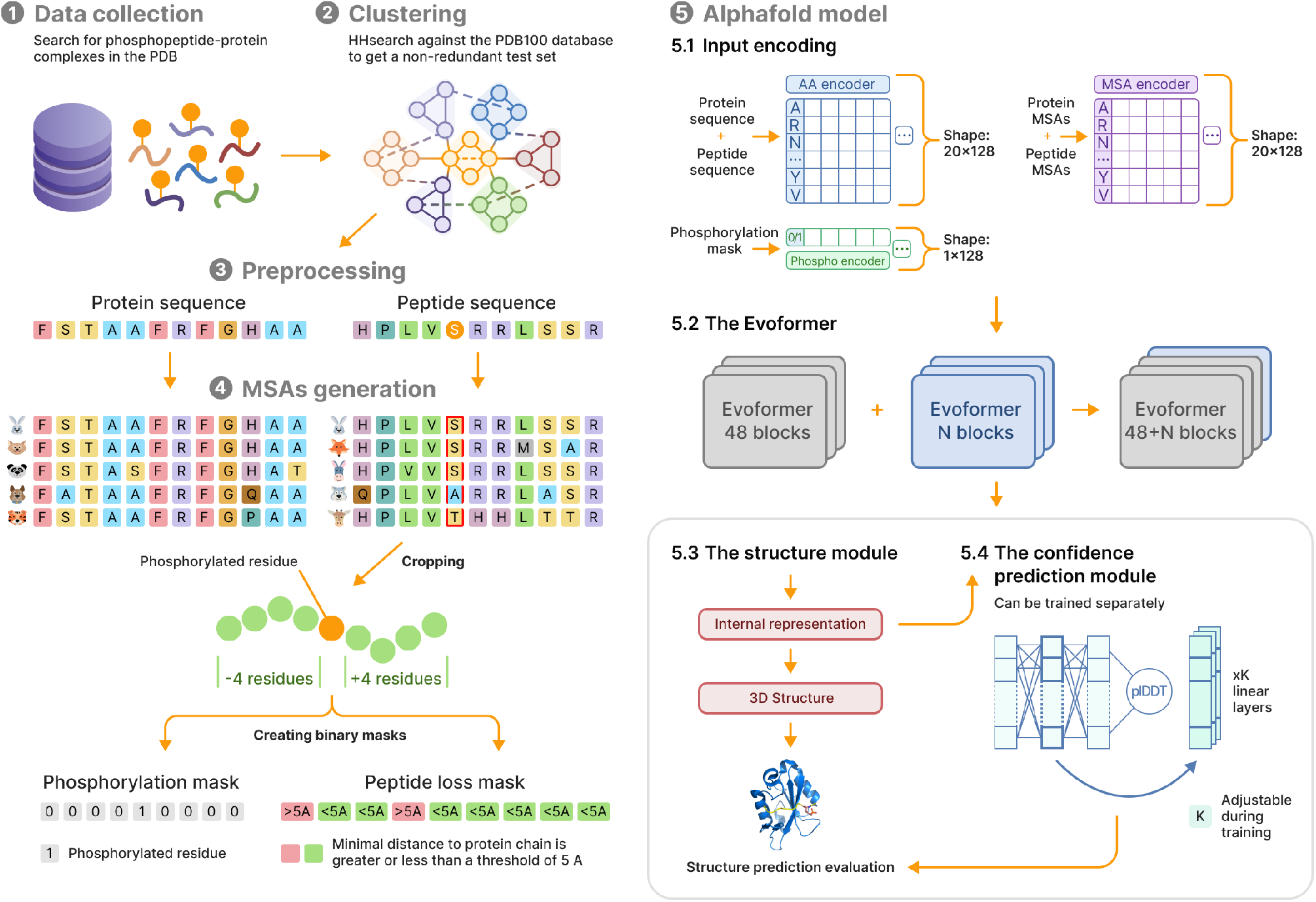
The overall pipeline of the Phospho-Tune model prediction process described in the paper. (1) We collect data by searching for and downloading known phosphopeptide-protein structure complexes from the Protein Data Bank (PDB); (2) We use HHsearch to generate hhr files, which are then used for grouping together proteins; (3) We create input features for the model by extracting information from the input file; (4) For input protein and peptide sequences, we generate Multiple Sequence Alignments (MSAs) derived from the Genetic Sequence Database. We crop peptide sequences to include only 4 residues before and after phosphorylated residue. We create binary masks, with the phosphorylation mask indicating the position of the phosphorylated residue within the sequence, and the peptide loss mask indicating peptide residues with a minimal distance to the protein chain below a threshold of 5 Å; (5.1) Input protein and peptide sequences, as well as MSA features, are transformed into latent representations, where each amino acid in the sequence is encoded into a 128-dimensional embedding vector. The encoding process is also applied to the phosphorylation mask; (5.2) The original AlphaFold model architecture consists of 48 Evoformer blocks, we introduce the option to increase this number by adding extra blocks, denoted as 48 + N blocks; (5.3) The Structure Module takes the final representations as input and generates a three-dimensional structure of phosphopeptide-protein complex. We evaluate the final predicted structure with RMSD, plDDT, and lDDT metrics; (5.4) We can train the PredictedLDDT module separately. This involves using the true per-residue lDDT score and the internal structure representation from the Structure Module as input. During training, we have the flexibility to adjust the number of linear layers.

**Fig. 6:**
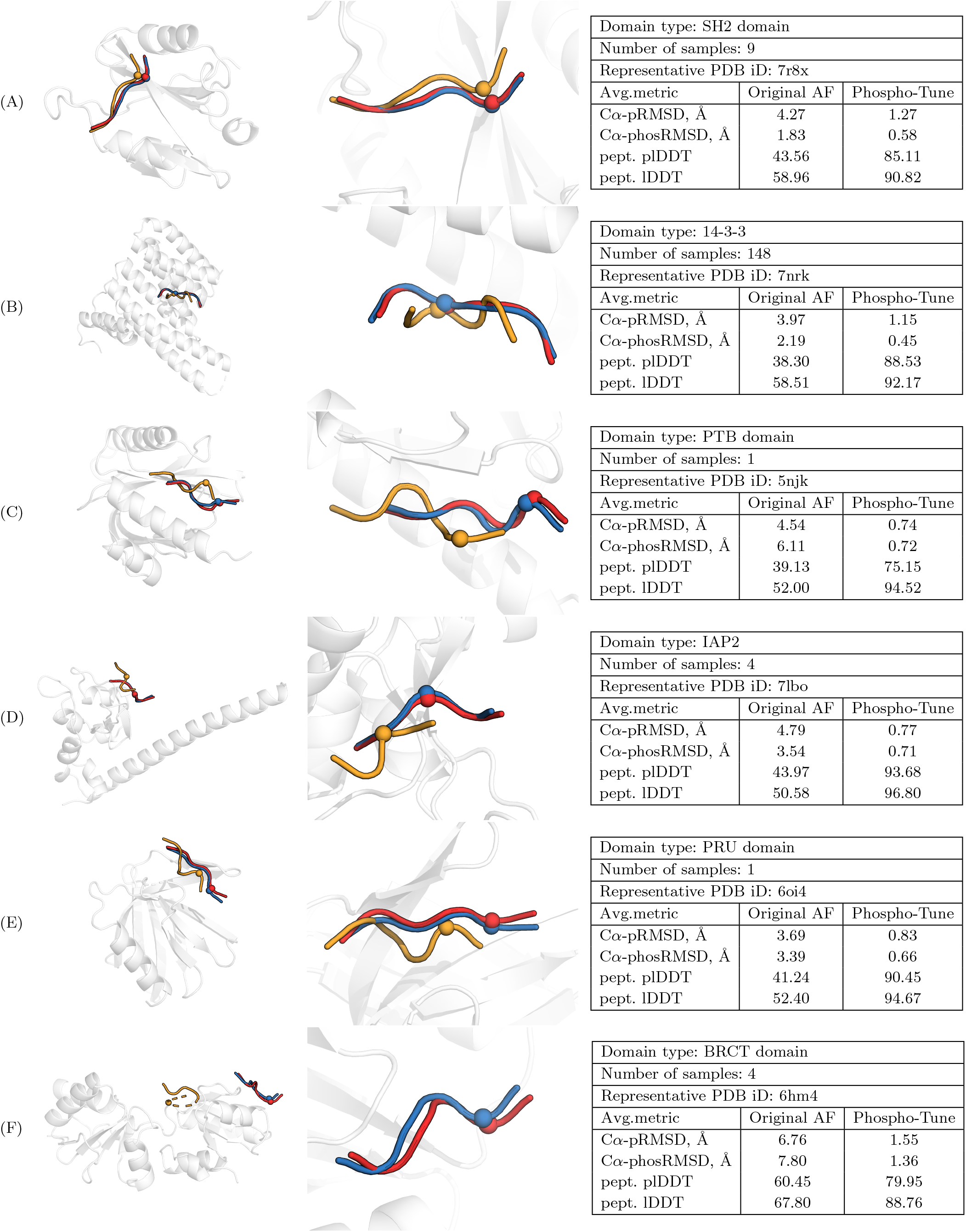
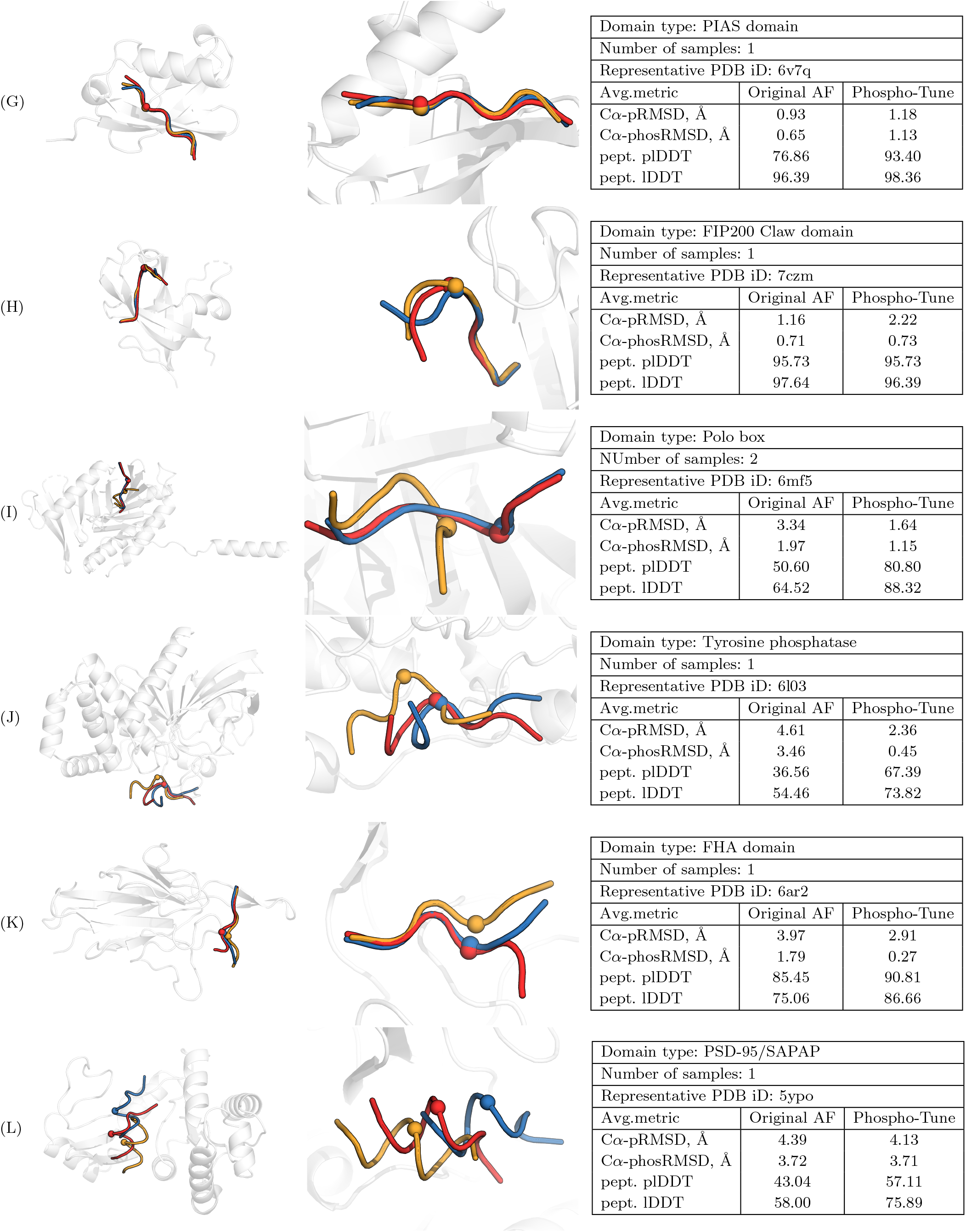
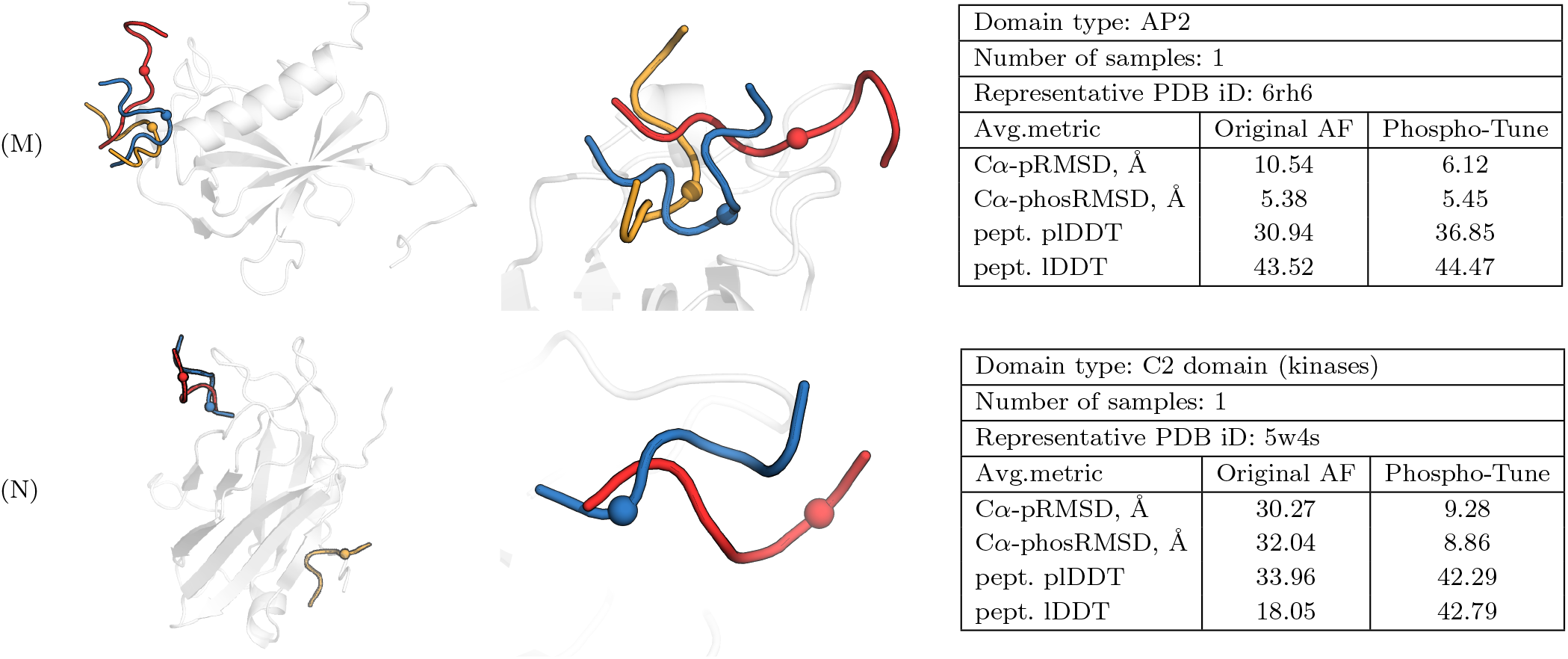
Comparative Visualization of Phosphopeptide Pose Prediction Accuracy. Each example is presented with two images: the left displays the overall protein structure with colored predictions for peptide structures, while the right provides a zoomed-in view of peptide alignments. In these visualizations, the true peptide structure is highlighted in red, the original AlphaFold model in yellow, and Phospho-Tune (fine-tuned AlphaFold) in blue. Additionally, each dot on the peptide structures signifies the CA atom position of a phosphorylated residue.

## Conclusion

In this study, we have introduced the Phospho-Tune model, a fine-tuned version of the AlphaFold model tailored for structural modeling of phosphopeptide-protein interactions across distinct protein families. Through a detailed fine-tuning process involving a specialized dataset and refined architecture, our study confirms the advantage of the Phospho-Tune model over the original AlphaFold. This is evidenced by enhanced accuracy and robust confidence scores, specifically tailored for precise evaluation of structural modeling, with a particular focus on phosphopeptides. This positions Phospho-Tune as a valuable tool for advancing our understanding of complex cellular processes and opening pathways for transformative progress in drug discovery and therapeutic research.

## Acknowledgments

This work was supported in part by the National Institutes of Health grants RM1135136, R01GM140098; by the National Science Foundation grants DMS-1664644, DMS-2054251. This research used resources of the Oak Ridge Leadership Computing Facility at the Oak Ridge National Laboratory, which is supported by the Office of Science of the U.S. Department of Energy.

